# Artificial light at night alters morphology, phenology, and reproductive capacity in an annual herb

**DOI:** 10.1101/2022.12.11.519667

**Authors:** Lu Xiao, Shuo Wang, Ayub M. O. Oduor, Zhihui Wang, Hongxiang Zhang, Yanjie Liu

**Affiliations:** State Key Laboratory of Black Soils Conservation and Utilization, Northeast Institute of Geography and Agroecology, Chinese Academy of Sciences, Changchun, 130102, China; Department of Applied Biology, Technical University of Kenya, P.O Box 52428-00200, Nairobi, Kenya; Shandong Key Laboratory of Eco-Environmental Science for Yellow River Delta, Shandong University of Aeronautics, Binzhou, Shandong, China; Heilongjiang Bayi Agricultural University, Daqing, 163319, China

**Keywords:** Artificial light at night, Phenological shifts, Light pollution, Plant traits, Ecological impact

## Abstract

The rapid global expansion of artificial light at night (ALAN) has spurred growing interest in its ecological impact on plant life. However, the effects of low-intensity ALAN on plants in their natural habitats remain largely unexplored, particularly concerning the influence of morphological and phenological responses on the overall fitness of these plants. We conducted a field experiment using the annual herbaceous plant *Elsholtzia densa* as a model species to assess the effects of ALAN on plant morphology, reproductive phenology, and reproductive capacity. The results indicate that ALAN increased the specific leaf area and elongated the top inflorescences, but it resulted in a reduction of secondary branches and a decrease in the proportion of individuals with undeveloped top inflorescences. Additionally, ALAN induced a shift in biomass allocation toward the aboveground parts of plants. It also accelerated the onset of budding, blooming, fruiting, and seed maturity by 3.4 to 6.2 days and caused a decrease in the number of fruity inflorescences. This study suggests that ALAN can significantly affect plant morphology, reproductive timing, and potentially the fitness of plants. While ALAN induces potentially adaptive changes in leaf area and biomass allocation, it may also disrupt plant-pollinator interactions and negatively impact plant reproductive capacity.

## 1. Introduction

To adapt to the cyclical changes in environmental light due to Earth’s rotation, a wide range of organisms, including algae, plants, and animals, have evolved molecular circadian clocks (Hölker et al. 2010; Yerushalmi and Green 2009). These clocks have reduced competition and increased coexistence among sympatric species by enabling organisms to optimize their physiological processes in synchrony with the diurnal cycle (Gaston et al. 2013; Krittika and Yadav 2019; Patke et al. 2020; Yerushalmi and Green 2009; Zhao et al. 2021). However, the natural darkness of Earth’s night, which has persisted for hundreds of billions of years, has been rapidly diminished in recent decades by the widespread use of artificial light at night (ALAN) across vast geographical areas (Falchi et al. 2016; Gaston 2018; Irwin 2018; Kyba et al. 2017). At a global scale, satellite data show that artificially lit outdoor areas expanded by 2.2% annually, with a 1.8% yearly increase in radiance intensity between 2012 and 2016 (Kyba et al. 2017). This rapid global increase of ALAN has led to light pollution, which has been shown to impact various species critically and potentially threaten ecosystems (Irwin 2018), making it a defining feature of the Anthropocene epoch. Consequently, there has been a growing interest in exploring the ecological effects of ALAN within the field of global change biology and urban ecology, particularly regarding its impacts on species behavior and ecosystem services (Davies and Smyth 2018; Hölker et al. 2021; Lockett et al. 2022).

A recent meta-analysis revealed that most studies on the impacts of altered photoperiod caused by ALAN have historically focused on animal behavior, physiology, and life history (Sanders et al. 2021). However, increasing attention is now being directed toward the effects of ALAN on plants. Numerous studies on urban trees, ornamental garden plant species, and crops have demonstrated that ALAN can significantly affect plant physiological processes, including photosynthesis, respiration, and stomatal conductance (Haque et al. 2015; Kim et al. 2015; Kwak et al. 2018; Kwak et al. 2017; Segrestin et al. 2021; Singhal et al. 2019). ALAN has also been found to influence plant morphology, affecting traits such as leaf size, stem elongation, and biomass allocation (Abonyo and Oduor 2024; Cieraad et al. 2023; Lockett et al. 2022). Moreover, ALAN can significantly impact plant phenology, particularly flowering time and plant-pollinator interactions (Abonyo and Oduor 2024; Bennie et al. 2018; Ffrench-Constant et al. 2016; Giavi et al. 2021; Liao et al. 2014; Meng et al. 2022). Despite these advances, the ecological consequences of ALAN on native flora in natural ecosystems remain understudied compared to urban and agricultural contexts. Therefore, significant knowledge gaps persist regarding the long-term effects of ALAN on plant community composition, biodiversity, and ecosystem functioning in natural habitats (Gaston et al. 2015; Secondi et al. 2019). Addressing these gaps should be a research priority to inform conservation strategies and mitigate ALAN’s potential negative impacts on natural ecosystems.

Given that plants rely on light as a crucial resource and signaling mechanism for regulating fundamental processes such as photoperiod responses, their growth and development are likely to be sensitive to ALAN (Bennie et al. 2016; Gaston et al. 2015; Singhal et al. 2019). Plants perceive light signals via specialised photoreceptors, including phytochromes, cryptochromes, and phototropins, which play pivotal roles in regulating growth and development (Quail 2002). The distinct spectral and temporal characteristics of ALAN may interfere with these photoreceptor-mediated pathways, disrupting the normal functioning of light-dependent processes (Heinen 2021; Meravi and Kumar Prajapati 2020). Furthermore, ALAN has the potential to disrupt circadian rhythms, which are essential for synchronising physiological processes with daily and seasonal light cycles. This disruption may alter the timing of gene expression and metabolic processes, leading to misalignment between the endogenous circadian clock and the external environment (Bordage et al. 2016; Fankhauser and Staiger 2002; Webb 2003). Such misalignment can have detrimental effects on plant growth, development, and fitness (Dodd et al. 2015; Pierik and de Wit 2014).

ALAN may also influence hormonal pathways that control plant growth and reproduction. For instance, Briggs (2006) hypothesized that low-intensity ALAN, which has a spectral composition different from direct sunlight, might be perceived by plants as a shaded environment. This perception can trigger shade avoidance responses, such as stem elongation and reduced branching, which are commonly mediated by enhanced auxin signaling and gibberellin production (Jiang et al. 2020; Pierik and de Wit 2014). These hormonal changes can lead to altered biomass allocation patterns, with plants investing more resources in vertical growth at the expense of leaf and root development (Hertel et al. 2021; Speißer et al. 2021). Recent studies have indeed shown that low-intensity ALAN, such as street lighting, can emit photosynthetically active radiation that affects the growth and resource allocation in vascular plants (Speißer et al. 2021) and algae (Diamantopoulou et al. 2021). While some plants may benefit from extended photoperiods under ALAN, others may experience reduced fitness due to the disruption of circadian rhythms and the metabolic costs associated with prolonged growth (Cathey and Campbell 1975; Gaston et al. 2013). The potential effects of ALAN on functional traits, particularly leaf function and resource allocation in natural plant populations, require further investigation to elucidate the underlying physiological mechanisms and their ecological consequences.

Understanding the impact of ALAN on plant flowering and other reproductive processes is critical, as ALAN-induced shifts in phenology may lead to mismatches between plants and their pollinators, thereby posing direct threats to the reproductive success of entomophilous plants. Research using satellite data (Zheng et al. 2021) and phenological observation data (Meng et al. 2022) suggests that ALAN might lead to an earlier start of the growing season. However, the specific effects of ALAN on flowering are variable, ranging from negative to neutral to positive (Abonyo and Oduor 2024; Bennie et al. 2018; Cathey and Campbell 1975), which highlights the complex and context-dependent nature of ALAN’s influence. Such phenological shifts can lead to pollinator mismatches, reducing pollination success and seed production, ultimately impacting dependent species (Giavi et al. 2021; Heinen 2021). ALAN may impact plant-pollinator networks by altering the timing of flowering and the availability of floral resources, which in turn can affect the abundance and diversity of pollinators (Knop et al. 2017; MacGregor et al. 2015). Furthermore, ALAN-induced changes in plant phenology and resource allocation may have cascading effects on plant-herbivore interactions (Bennie et al. 2015; Grenis and Murphy 2019) and plant-plant interactions, such as competition (Abonyo and Oduor 2024).These alterations in species interactions can potentially reshape the structure and composition of plant communities (Bennie et al. 2016; Bennie et al. 2018; Hölker et al. 2021; Speißer et al. 2021). Given the close link between reproductive traits and fitness (Violle et al. 2007) , ALAN’s capacity to alter these traits could directly affect the viability of natural plant populations. Therefore, to fully understand the ecological consequences of ALAN on plant reproduction in the wild, research should prioritize field experiments over laboratory or greenhouse studies, allowing for a more accurate assessment of plant responses in their natural environments and within the context of their ecological communities (Secondi et al. 2019).

This study investigated the effects of artificial light at night (ALAN) on the fitness of the widespread annual herb *Elsholtzia densa* Benth., with a particular focus on morphological traits, reproductive phenology, and reproductive capacity of the species. We tested the following predictions: (1) ALAN increases specific leaf area of plant; (2) ALAN enhances plant height and total biomass; (3) ALAN advances the onset of budding, blooming, and fruiting as a result of extended photoperiod and disruption of circadian rhythms; and (4) ALAN increases both the size and number of inflorescences due to changes in vegetative growth and reproductive investment. These hypotheses reflect the increasing recognition that ALAN can interfere with light signaling pathways, alter photoperiod-sensitive processes, and affect internal resource distribution in plants. By examining these responses in a natural setting, the study aims to elucidate the mechanisms by which ALAN influences plant performance, thereby contributing to a more comprehensive understanding of its ecological impacts on plant fitness and population dynamics.

## 2. Materials And Methods

### 2.1. Study sites and species

This study is part of an ongoing long-term ALAN manipulation experiment conducted in a temperate continental monsoon climate grassland in Inner Mongolia, China (50.17°N, 119.39°E, 523 m a.s.l.). This grassland experiences a mean annual precipitation of 375 mm and a mean annual temperature of –3 °C (Liu et al. 2021) . The broader experiment encompasses 40 square plots, each 4 m², arranged in four combinations of two light pollution levels (ALAN vs. no-ALAN) and two animal-exclusion levels (open cage vs. closed cage), with ten replicates for each combination. The present study specifically utilized 20 plots from the larger experiment, employing only the no-animal-exclusion treatment (open cage) and the two light pollution levels.

To ensure homogeneity in environmental conditions across all plots, we selected a relatively uniform area within the grassland, characterized by similar soil texture, moisture levels, and nutrient availability. The soil was initially tilled, and clonal propagules were removed to homogenize species composition. On April 18, 2021, artificial communities were re-established using seeds from 54 common herbaceous species collected in 2020 from various local grassland sites, ensuring a diverse representation of populations and genetic diversity. The seeds were mixed and randomly distributed across all plots, maintaining a consistent species proportion. All species included were capable of independent propagation without human intervention. We chose the widespread annual herb, *E. densa*, as our model organism to assess the impacts of ALAN on plant morphology, reproductive phenology, and reproductive capacity. Its dominance in our experimental plots, where it exhibited a relative cover of 33.78 ± 3.0% in 2021 (Supplementary Data Fig. S1), alongside its broad distribution from East Afghanistan to Southeast Siberia and China (POWO database; https://powo.science.kew.org), made it an ideal candidate for studying the effects of light pollution on common plant species.

### 2.2. Set up of the artificial-light-at-night experiment

The ALAN treatment was implemented using LED spotlights (10 W, DC 12–24 V, IP 66, cool white 6500 K; Shenzhen Mingkeming Lighting Co., Ltd., China) mounted on metal frames 2 meters above the center of each plot. These lights emit photosynthetically active radiation (PAR; Supplementary Data Fig. S2). To minimize lateral light radiation and prevent interference between plots, each spotlight was encased in a cubic lampshade (30 × 30 × 30 cm) made of stainless steel, and plots were spaced 2-3 meters apart. To replicate the intensity of light pollution typically experienced at ground level, we covered each lampshade with two layers of white nylon netting and one layer of black light-shading netting. This setup resulted in a consistent light intensity of 21.15 ± 0.30 lx, comparable to that found under streetlights (Bennie et al. 2016; Speißer et al. 2021). This value represents the spatiotemporal average per plot, based on measurements taken at five randomly selected points on the ground within each plot. While some variability existed, statistical analyses confirmed that light intensity remained significantly different between ALAN and No-ALAN treatments. The lamps for the ALAN treatment were regulated by a photoelectric switch (XT KG-F, Zhejiang Xintuo New Energy Co., Ltd., China), which automatically turned the lights on when ambient light was below 30 lux and off when above this threshold during the growing season. Control plots (no-ALAN) also had metal frames and lampshades made of stainless steel covered with nylon netting but lacked LED spotlights. Light intensity measurements in control plots using a SpectraPen LM500 spectroradiometer (Photon Systems Instruments) yielded zero readings, indicating levels below the device’s detection limit (< 0.1 lux).

#### 2.3. Trait measurements

We measured 14 traits related to morphology, reproductive phenology, and reproductive capacity of *E. densa* from the onset of budding on July 6, 2021, to the first frost on September 23, 2021 (Table 1). Ten of these traits were measured non-destructively on 10 randomly selected and marked individuals of *E. densa* within a 1 × 1 m subplot at the center of each plot. From July 6 to September 23, we monitored the marked *E. densa* individuals daily to record the onset of budding, blooming, fruiting, and seed maturity. The onset of budding was defined as the appearance of the first bud, blooming as the emergence of the first purple inflorescence, fruiting as the initial swelling of the inflorescence, and seed maturity as the presence of the first dehiscent inflorescence, following the methodology of Dorji et al. (2013). At the end of the growing season (September 23), we measured branch number and reproductive height of each marked individual plant. Branches were classified as primary if arising directly from the main stem, and as secondary if originating from the primary branches, like the approach used by Li et al. (2016). To assess the impact of ALAN on reproductive capacity, we recorded the number of fruiting inflorescences (inflorescences bearing seeds) per individual and the number of individuals with undeveloped top inflorescences (remaining at the bud stage until the end of the growing season). In this study, we used the number of fruiting inflorescences as a proxy for reproductive capacity in *E. densa* due to the challenges associated with directly measuring seed production in the field. Premature seed fall makes it difficult to accurately quantify the total number of seeds produced by individual plants. The number of fruiting inflorescences, however, is a more reliable and practical measure that can be assessed in the field without the risk of underestimating reproductive output due to seed loss. The number of fruiting inflorescences often correlates with seed production (Kudo and Harder 2005; Rodger et al. 2021), serving as a proxy for reproductive capacity and offering insights into the impact of ALAN on plant fitness. This approach is supported by studies showing that fruiting inflorescences can be a reliable indicator of reproductive capacity, especially when direct seed production is challenging to measure (Kudo and Harder 2005; Waser and Ollerton 2006). Additionally, noting a potential difference in top inflorescence length between ALAN and no-ALAN plots (Fig. S3), we measured this trait and calculated the proportion of individuals with undeveloped top inflorescences in each plot as: the number of individuals with undeveloped top inflorescence / 10 marked individuals per plot.

**Table 1.** The 14 traits measured in this study.

| Category | Trait | Measuring time | Replication No. |
| --- | --- | --- | --- |
| Morphology | *Specific leaf area* | 27 August, 2021 | 20 |
|  | *Primary branch number* | 23 September, 2021 | 200 |
|  | *Secondary branch number* | 23 September, 2021 | 200 |
|  | *Aboveground biomass fraction* | 27 August, 2021 | 97 |
|  | *Height* | 23 September, 2021 | 200 |
|  | *Total biomass* | 27 August, 2021 | 97 |
| Reproductive phenology | *Budding date* | From 6 July to | 200 |
|  | *Blooming date* | 23 September, 2021 | 200 |
|  | *Fruiting date* | (before plant budding | 200 |
|  | *Seed maturity date* | to the start of frost) | 200 |
| Reproduction capacity | *Length of top inflorescence* | 23 September, 2021 | 200 |
|  | *Proportion of individuals with undeveloped top inflorescence* | 23 September, 2021 | 20 |
|  | *Inflorescence biomass fraction* | 27 August, 2021 | 97 |
|  | *Fruity inflorescence number* | 23 September, 2021 | 200 |

We conducted destructive sampling to measure four additional traits. To assess the response of specific leaf area to ALAN, we collected ten mature, fully expanded, and healthy leaves from five unmarked *E. densa* individuals adjacent to the central subplot within each plot on August 27. Each individual contributed two leaves. We scanned the entire leaves using a CanoScan LiDE 300 electronic scanner (Canon Inc., Lake Success, NY, USA) and calculated total leaf area with ImageJ software (version 1.53g). Subsequently, the leaves were dried at 60 °C for 72 hours and weighed. Specific leaf area was calculated for each plot as the total leaf area divided by the total dry leaf biomass of the corresponding plot. On the same day (August 27), we also measured the aboveground, inflorescence, and total biomass of *E. densa* under both ALAN treatments. For this, we randomly selected an additional 3-6 individuals adjacent to each subplot, resulting in 45 individuals for ALAN plots and 52 for no-ALAN plots. We harvested the belowground parts, inflorescences, and other aboveground parts separately for each of the 97 individuals. All materials were dried at 60°C for 72 hours and weighed. From this data, we calculated the total biomass (belowground parts + inflorescences + other aboveground parts), aboveground biomass fraction (sum of inflorescences and other aboveground parts divided by total biomass), and inflorescence biomass fraction (inflorescences divided by total biomass) for each *E. densa* individual. It is important to note that both leaf function and biomass allocation measurements were taken during the vigorous growth stage (August 27), when some inflorescences were blooming but not yet senescing. This timing was chosen to capture trait values representative of the plant’s peak physiological performance. As demonstrated in previous studies, the timing of trait measurements can substantially influence observed patterns and their responses to environmental conditions (Famiglietti et al. 2024). Given the extended reproductive period of *E. densa* (approximately 50 days), the relatively small ALAN-induced reproductive phenology shift (3-6 days) likely had only minimal impact on the observed differences in leaf functionality and biomass allocation. Sampling too early could have missed important developmental effects of ALAN on plant traits, while sampling too late might have introduced confounding effects due to senescence-related trait decline.

### 2.4. Statistical analyses

To investigate the effects of ALAN on various plant traits, we fitted a series of Bayesian multilevel models using the *brm* function from the *brms* package (Bürkner 2017) in R v.4.0.2 (R Core Team 2020). These models accounted for the hierarchical structure of the data and the non-independence of repeated measurements. A total of 14 traits were analyzed, with ALAN included as a fixed effect and the specific trait as the response variable in each model. Prior to selecting the appropriate model for each trait, we conducted exploratory data analysis to assess the distribution pattern. Histograms were used to visually inspect the distribution of each variable. The number of fruity inflorescences, primary branches, and secondary branches were modeled using negative binomial distributions. The proportion of individuals with undeveloped top inflorescences was modeled using a binomial distribution. Plant height and total biomass were modeled with gamma distributions, while the above-ground and inflorescence biomass fractions were modeled using beta distributions. The remaining traits, including specific leaf area (square root transformed), budding date, blooming date, fruiting date, seed maturity date, and length of top inflorescence were modeled using Gaussian distributions. To account for the non-independence of individuals within plots, plot identity was included as a random effect in all models, except for those analyzing specific leaf area and the proportion of individuals with undeveloped top inflorescences, which were measured at the plot level. All models used the default priors provided by the *brms* package and were run with four independent chains, each consisting of 6000 iterations, including 2000 warm-up iterations. Model fit was assessed using the *pp_check* function from *brms*, which compares model-generated data with the observed data via visual inspection of density and scatter plots (Gabry et al. 2019). A model was considered adequate if the simulated data closely approximated the observed data in terms of shape and spread. To select the optimal model for each trait, we used the *loo* function from the *brms* package, which implements leave-one-out cross-validation (LOO-CV) to estimate the expected log predictive density and the Widely Applicable Information Criterion (WAIC) (Vehtari et al. 2017). The model with the lowest WAIC was selected as the optimal model for each trait, indicating the best balance between model fit and complexity. The significance of the ALAN treatment was assessed based on the credible intervals of the posterior distribution. Specifically, the ALAN treatment was deemed significant when the 95% credible interval of the posterior distribution did not include zero, and marginally significant if the 90% credible interval did not overlap zero.

## 3. Results

### 3.1. Effects of ALAN on morphological traits

Two of the six morphological traits of *E. densa* were significantly affected by ALAN (Table 2). Specifically, ALAN significantly increased the specific leaf area (+9.2%; Fig. 2a, Table 2) but decreased the number of secondary branches (−44.3%; Fig. 2c, Table 2) compared to the no-ALAN treatment. Relative to the no-ALAN treatment, ALAN also tended to increase the aboveground biomass fraction (+0.68%; Fig. 2f, Table 2). The number of primary branches, height, and total biomass were not significantly affected by ALAN treatment (Fig. 2b, d, e; Table 2).

**Fig. 1.**
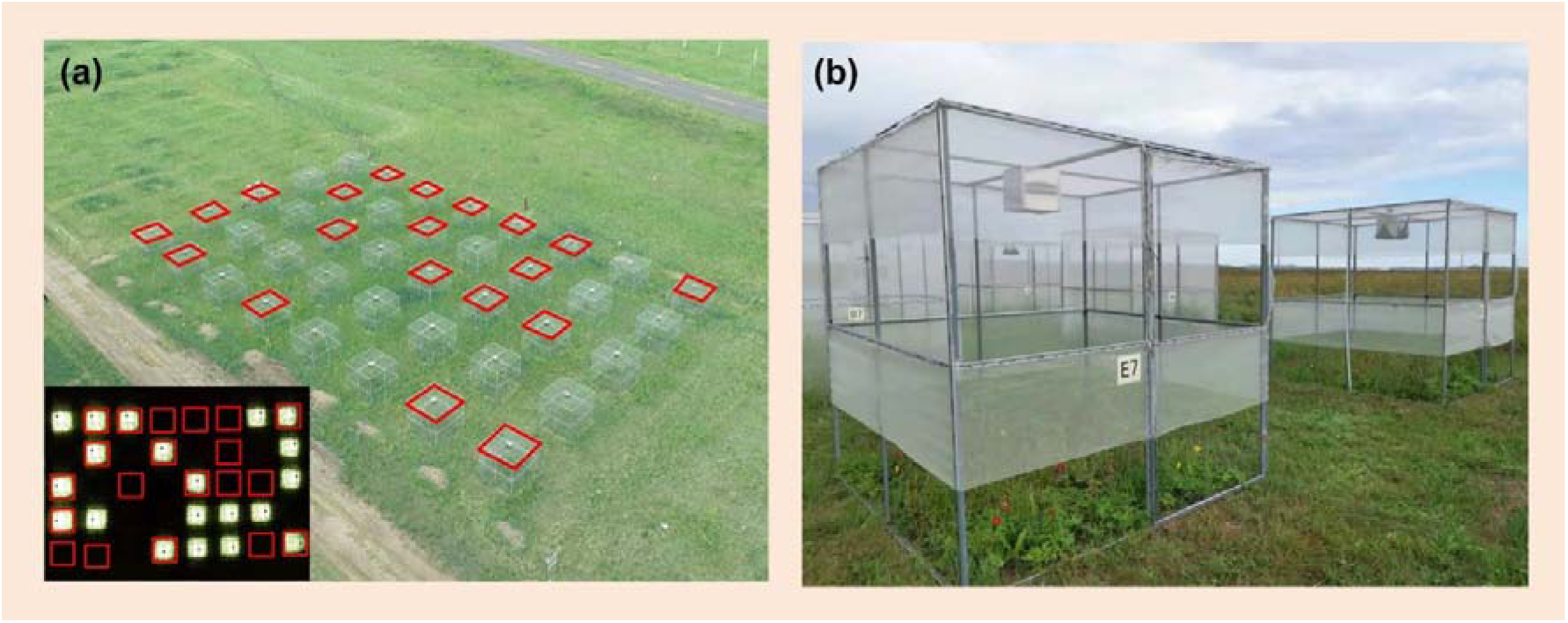
(a) Top view of the entire experimental setup, showcasing the 20 open cages (no animal exclusion) used in this study (marked with borders), both during the day and at night (lower left inset). (b) Close-up photograph of a representative open cage plot.

**Fig. 2.**
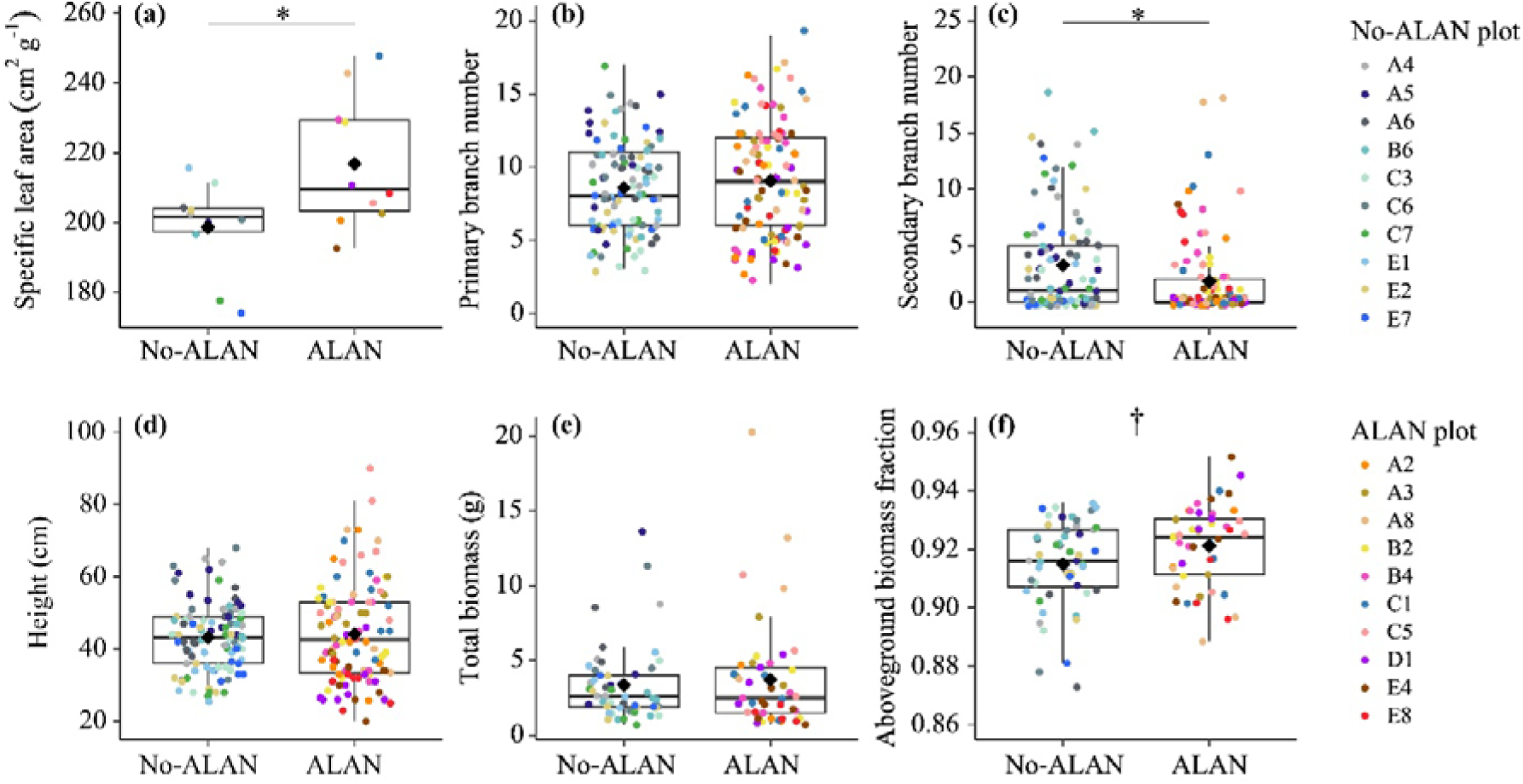
Effects of artificial light at night (ALAN) on specific leaf area (a), primary branch number (b), secondary branch number (c), plant height (d), total biomass (e), and aboveground biomass fraction (f). Asterisks (*) indicate that the 95% credible intervals of the estimated effects of ALAN do not include zero, indicating a statistically significant difference. Daggers (†) represent 90% credible intervals that do not include zero, suggesting a marginally significant difference. Details on each trait are provided in Table 2.

**Table 2.** Output of Bayesian models testing the effects of artificial light at night (ALAN) on each of the 14 traits observed in this study. Asterisks (*) and daggers (†) indicate terms whose 95% and 90% credible intervals (CIs) do not overlap with zero, respectively.

| No. | Trait | brm family | Term | Fixed effect |  |  | Random effect |
| --- | --- | --- | --- | --- | --- | --- | --- |
|  |  |  |  | Estimate | 95% CIs | 90% CIs | Estimate (95% CIs) |
| 1 | *Specific leaf area*<br>(*n* = 20) | Gaussian | Intercept (No-ALAN) * | 14.087 | (13.707, 14.473) | (13.767, 14.404) | — |
|  |  |  | ALAN * | 0.630 | (0.067, 1.181) | (0.171, 1.084) | — |
| 2 | *Primary branch number*<br>(*n* = 200) | Negative binomial | Intercept (No-ALAN) * | 2.132 | (2.009, 2.252) | (2.03, 2.232) | 0.14 (0.04, 0.25) |
|  |  |  | ALAN | 0.060 | (-0.110, 0.237) | (-0.08, 0.203) | — |
| 3 | *Secondary branch number*<br>(*n* = 200) | Negative binomial | Intercept (No-ALAN) * | 1.192 | (0.763, 1.659) | (0.832, 1.575) | 0.26 (0.01, 0.78) |
|  |  |  | ALAN * | -0.637 | (-1.322, -0.006) | (-1.183, -0.111) | — |
| 4 | *Aboveground biomass fraction*<br>(*n* = 97) | Beta | Intercept (No-ALAN) * | 2.375 | (2.32, 2.44) | (2.326, 2.425) | — |
|  |  |  | ALAN † | 0.081 | (-0.007, 0.170) | (0.007, 0.155) | — |
| 5 | *Height*<br>(*n* = 200) | Gamma | Intercept (No-ALAN) * | 0.024 | (0.021, 0.027) | (0.021, 0.026) | 0.01 (0.00, 0.02) |
|  |  |  | ALAN | 0.001 | (-0.004, 0.004) | (-0.003, 0.003) | — |
| 6 | *Total biomass*<br>(*n* = 97) | Gamma | Intercept (No-ALAN) * | 0.330 | (0.234, 0.438) | (0.25, 0.418) | 0.13 (0.08, 0.21) |
|  |  |  | ALAN | 0.006 | (-0.139, 0.153) | (-0.114, 0.124) | — |
| 7 | *Budding date*<br>(*n* = 200) | Gaussian | Intercept (No-ALAN) * | 205.513 | (204.369, 206.629) | (204.575, 206.446) | 0.49 (0.02, 1.35) |
|  |  |  | ALAN * | -3.588 | (-4.954, -2.243) | (-4.713, -2.464) | — |
| 8 | *Blooming date*<br>(*n* = 200) | Gaussian | Intercept (No-ALAN) * | 229.190 | (228.101, 230.243) | (228.313, 230.059) | 1.29 (0.39, 2.46) |
|  |  |  | ALAN * | -5.494 | (-7.673, -3.457) | (-7.294, -3.785) | — |
| 9 | *Fruiting date*<br>(*n* = 200) | Gaussian | Intercept (No-ALAN) * | 238.063 | (237.033, 239.111) | (237.216, 238.896) | 0.96 (0.09, 2.07) |
|  |  |  | ALAN * | -3.335 | (-5.107, -1.583) | (-4.814, -1.885) | — |
| 10 | *Seed maturity date*<br>(*n* = 200) | Gaussian | Intercept (No-ALAN) * | 255.622 | (254.520, 256.745) | (254.722, 256.548) | 0.81 (0.04, 2.12) |
|  |  |  | ALAN * | -4.526 | (-6.609, -2.462) | (-6.259, -2.784) | — |
| 11 | *Length of top inflorescence*<br>(*n* = 200) | Gaussian | Intercept (No-ALAN) * | 5.373 | (4.942, 5.798) | (5.015, 5.728) | 0.23 (0.01, 0.60) |
|  |  |  | ALAN * | 0.836 | (0.255, 1.429) | (0.348, 1.329) | — |
| 12 | *Proportion of individuals with undeveloped top inflorescence*<br>(*n* = 20) | Binomial | Intercept (No-ALAN) * | -2.105 | (-2.775, -1.509) | (-2.658, -1.598) | — |
|  |  |  | ALAN * | -1.920 | (-3.786, -0.480) | (-3.421, -0.693) | — |
| 13 | *Inflorescence biomass fraction*<br>(*n* = 97) | Beta | Intercept (No-ALAN) * | -1.809 | (-2.015, -1.610) | (-1.977, -1.645) | 0.22 (0.04, 0.41) |
|  |  |  | ALAN | 0.020 | (-0.264, 0.299) | (-0.215, 0.254) | — |
| 14 | *Fruity inflorescence number*<br>(*n* = 200) | Negative binomial | Intercept (No-ALAN) * | 3.208 | (2.966, 3.446) | (3.007, 3.404) | 0.32 (0.17, 0.51) |
|  |  |  | ALAN † | -0.285 | (-0.628, 0.050) | (-0.567, -0.008) | — |

### 3.2. Effects of ALAN on reproductive phenology

ALAN significantly affected the reproductive phenology of *E. densa* (Table 2). Specifically, ALAN advanced the averaged Julian days for the onset of budding, blooming, fruiting, and seed maturity by 3.4, 6.2, 3.4, and 4.6 days, respectively (Fig. 3, Table 2).

**Fig. 3.**
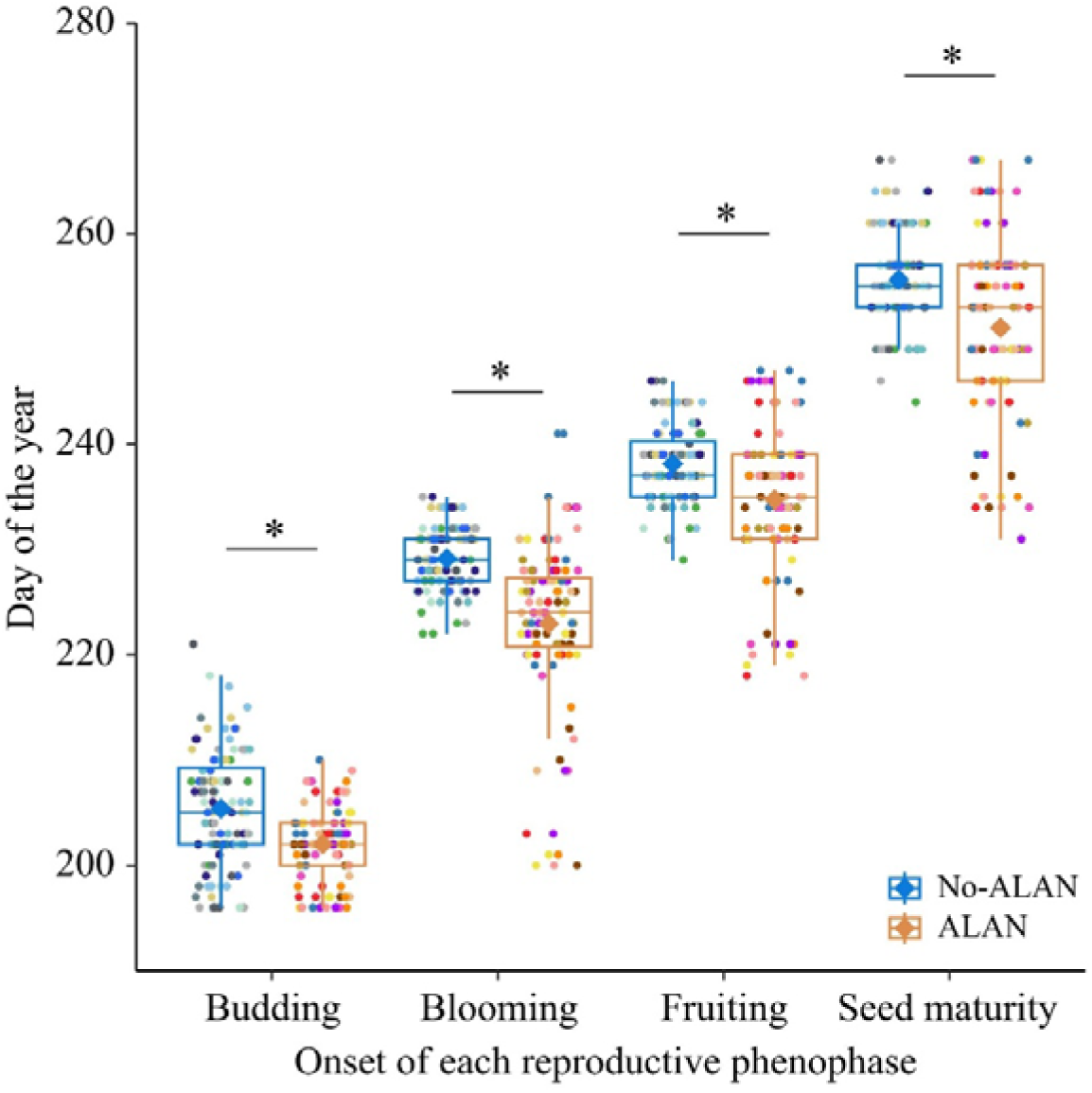
Effects of artificial light at night (ALAN) on the onset of different reproductive phenophases (budding, blooming, fruiting, and seed maturity), expressed as Julian days for the year 2021. Asterisks (*) indicate that the 95% credible intervals of the estimated effects of ALAN do not include zero, indicating a statistically significant difference. Details on each trait are provided in Table 2. Data points in different colors represent measurements from various plots, as detailed in Figure 2.

### 3.3. Effects of ALAN on reproductive capacity

ALAN significantly increased the length of the top inflorescences of *E. densa* (+29.2%; Fig. 4a, Table 2). Additionally, ALAN significantly decreased the proportion of individuals with undeveloped top inflorescences (−81.8%; Fig. 4b, Table 2). ALAN caused a marginally significant reduction in the number of fruity inflorescences relative to the no-ALAN treatment (−25.2%; Fig. 4d, Table 2). The ALAN treatment did not have a significant effect on the biomass fraction of inflorescence (Table 2).

**Fig. 4.**
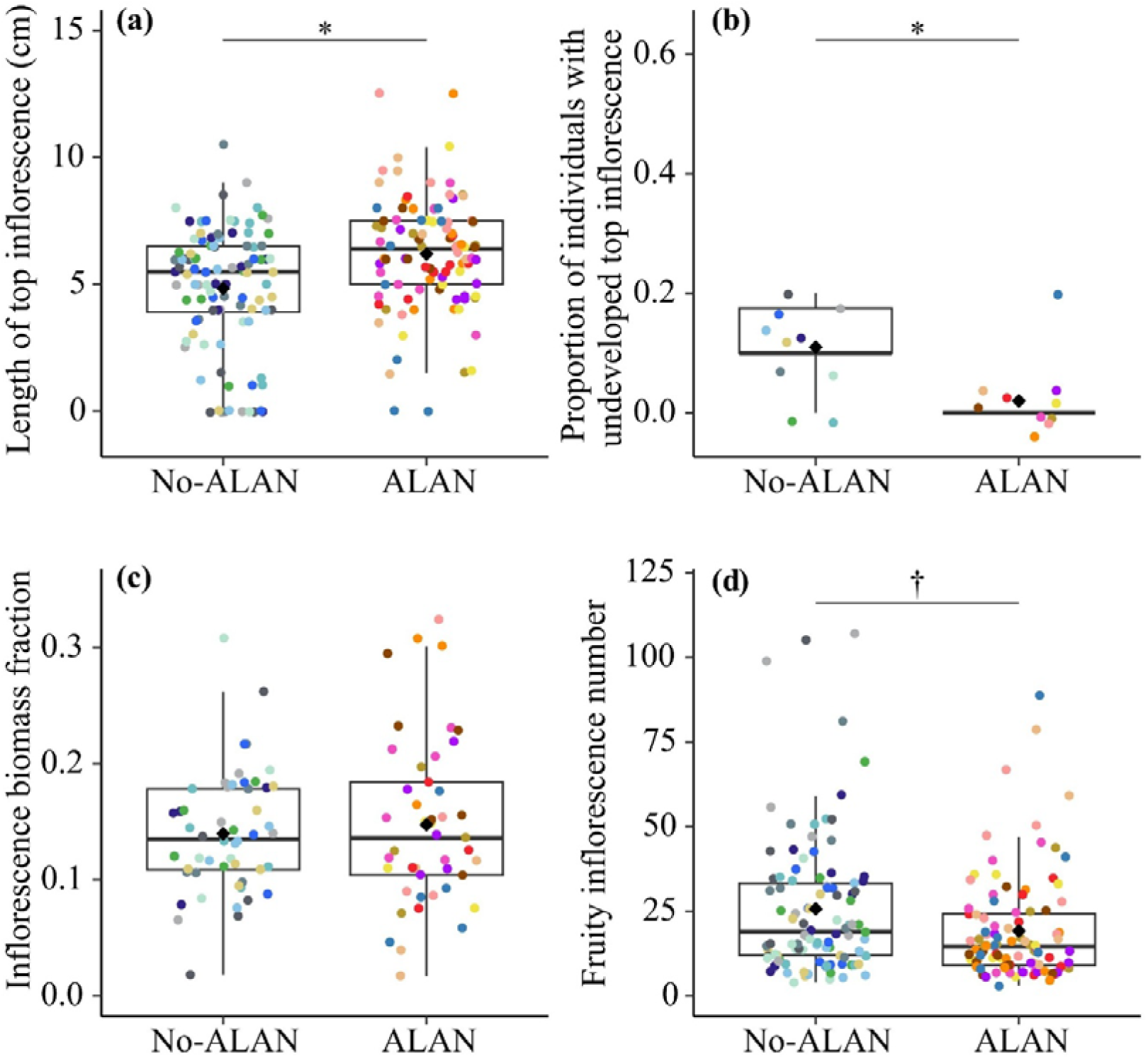
Effects of artificial light at night (ALAN) on length of top inflorescence (a), proportion of individuals with undeveloped top inflorescence (b), inflorescence biomass fraction (c), and fruity inflorescence number (d). Asterisks (*) indicate that the 95% credible intervals of the estimated effects of ALAN do not include zero, indicating a statistically significant difference. Daggers (†) represent 90% credible intervals that do not include zero, suggesting a marginally significant difference. Information on each trait can be found in Table 2. Data points in different colors represent measurements from various plots, as detailed in Fig. 2.

## 4. Discussion

Our study contributes to the growing body of literature on the ecological effects of ALAN on plants (Bennie et al. 2016; Gaston et al. 2017) by demonstrating that ALAN significantly influenced the morphology, reproductive phenology, and reproductive capacity of natural populations of the annual herb *E. densa*. Specifically, ALAN increased specific leaf area and aboveground biomass, while decreasing secondary branch numbers, suggesting a shift in resource allocation towards vertical growth and light capture. Additionally, ALAN advanced the timing of key reproductive stages, including budding, blooming, fruiting, and seed maturity, indicating a disruption in the plant’s natural phenological cycles. These findings highlight the multifaceted effects of ALAN on plant development and reproduction.

The morphological responses of *E. densa* to ALAN present a complex picture, possibly reflecting a combination of adaptive strategies. The observed increase in specific leaf area under ALAN conditions aligns with previous research demonstrating that high light exposure promotes leaf expansion and reduced leaf thickness to maximize light capture and photosynthetic efficiency (Poorter et al. 2009). This adaptation, coupled with the observed tendency for increased aboveground biomass allocation under ALAN, suggests that *E. densa* may be strategically prioritizing resources towards leaves and stems at the expense of belowground structures. This shift in resource allocation likely optimizes light capture and photosynthetic performance under extended photoperiod caused by ALAN. Similar shifts in biomass allocation in response to ALAN have been documented in other plant species (Abonyo and Oduor 2024). The reduction in secondary branches, a shade-avoidance response, might also prioritize vertical growth and leaf expansion for maximizing light interception (Gratani 2014). However, the lack of significant effects of ALAN on the number of primary branches, height, and total biomass highlights the importance of considering multiple morphological traits when assessing the ecological consequences of ALAN. While these morphological changes could potentially enhance the competitive ability of *E. densa* by optimizing light capture, they may also make the plant more susceptible to environmental stressors such as drought or herbivory. This is because a greater allocation of resources to above-ground structures might reduce the plant’s resilience to such stressors (Gupta et al. 2020). These findings suggest that while ALAN-induced changes in resource allocation may provide short-term benefits in terms of light capture, they could have long-term consequences for plant survival and fitness in natural ecosystems. Future research should investigate whether these morphological and phenological changes translate into increased fitness and survival of *E. densa* in natural ecosystems, or if they have unintended consequences for the plant’s overall ecological performance.

The observed advancement in the budding, blooming, fruiting, and seed maturity of *E. densa* under ALAN condition are consistent with studies showing that ALAN can cause earlier leaf-out and flowering in plants, extend the growing season, and alter reproductive timing (Cieraad et al. 2023; Ffrench-Constant et al. 2016; Meng et al. 2022). This acceleration of reproductive phenology is likely due to the extended photoperiod and increased light availability under ALAN, which modifies the plants’ perception of seasonal cues and triggers earlier initiation of reproductive processes (Gaston et al. 2017). Our observations suggest that *E. densa* is a long-day plant, typically initiating bud differentiation in mid-July under ambient light when day length is approximately 15.8 hours. The light spectrum emitted by our LED spotlights, containing red and far-red light (650nm-740nm, Fig. S2), can activate phytochromes, key light-sensing molecules that influence flowering (Bennie et al. 2016). This suggests that ALAN likely induces *E. densa* to perceive a shorter night length, triggering earlier flowering. This early flowering likely extends to subsequent phenological stages, as evidenced by the synchronized advancement across blooming, fruiting, and seed maturity observed in our study. While ALAN is primarily associated with photoperiod extension, other mechanisms, such as hormonal regulation, may also play a role. For instance, gibberellins, which promote cell elongation and stem growth, may increase under ALAN conditions and contribute to the observed phenological shifts (Sena et al. 2024). Similarly, abscisic acid, involved in stress responses and dormancy, may also be affected, potentially altering the timing of leaf senescence (Lazzarin et al. 2021). Likewise, jasmonates, which regulate defense and reproductive development, may be disrupted by ALAN, influencing flowering and fruit maturation (Lazzarin et al. 2021). Therefore, it is plausible that ALAN-induced hormonal changes, alongside altered photoperiod perception, contribute to the phenological shifts observed in *E. densa*.

The phenological shifts observed in *E. densa* under ALAN may have important ecological consequences, particularly by disrupting the timing of interactions with pollinators, seed dispersers, and other species (Hölker et al. 2010; Knop et al. 2017). Although the observed shift in flowering time, ranging from 3 to 6 days in our study, may appear small relative to the total flowering period (*ca*. 65 days), even slight mismatches can affect ecological interactions. Diurnal pollinators such as bees and butterflies may miss peak floral resource availability (Giavi et al. 2021), while nocturnal pollinators like moths and bats may reduce activity under continuous illumination (Knop et al. 2017). Similarly, earlier seed maturation may become decoupled from the peak activity of seed dispersers, potentially lowering dispersal efficiency and recruitment success (Schupp et al. 2010). Given that the ecological consequences of ALAN are complex and can extend beyond direct effects on individual plant species (Giavi et al. 2021; Macgregor et al. 2017), there is a need for more comprehensive studies to fully understand the ecological consequences of ALAN.

The present study reveals the complex effects of ALAN on plant reproductive traits, demonstrating both potential benefits and drawbacks (Fig. 4). ALAN significantly increased the length of top inflorescences (+29.2%) and decreased the proportion of individuals with undeveloped top inflorescences (-81.8%), suggesting a stimulatory effect on inflorescence growth and maturation. These responses may be driven by extended photoperiod and increased light availability, which are known to enhance vegetative growth and promote the development of reproductive structures (Abonyo and Oduor 2024; Ffrench-Constant et al. 2016). However, the marginally significant reduction in the number of fruiting inflorescences under ALAN (−25.2%) raises the possibility of trade-offs or constraints on reproductive success. The lack of change in inflorescence biomass fraction suggests that ALAN influences the timing and developmental efficiency of reproduction rather than altering overall reproductive investment. Additionally, ALAN may affect plant physiology and resource allocation, contributing to the observed shifts in reproductive traits (Heinen 2021). To fully understand the implications of these changes, future research should investigate how ALAN affects broader ecological interactions, including plant-pollinator dynamics, herbivory, and competition within plant communities (Heinen 2021; Knop et al. 2017) . Such studies would provide a more holistic understanding of ALAN’s multifaceted impacts on plant reproductive success.

Changes in the physical characteristics and reproductive timing of *E. densa* due to ALAN may lead to reduced fitness. While directly measuring seed production in the field is difficult because of early seed drop, we used the number of fruiting inflorescences as a proxy for reproductive capacity and fitness. Our results show that ALAN led to fewer fruiting inflorescences and a lower proportion of plants with undeveloped top inflorescences. Although our additional study (data not shown) did not find ALAN exposure in the parent generation to affect seed germination or seedling survival, the overall decrease in reproductive capacity suggests a potential negative impact on *E. densa* fitness. However, the long-term consequences for future generations remain uncertain. Importantly, the significant reduction in inflorescences of this nectar-rich plant could lead to food shortages for pollinating insects like bees, potentially disrupting ecosystem services through a bottom-up effect (Jiang et al. 1996)

## 5. Conclusions

In conclusion, our study contributes to the growing body of literature on the ecological effects of ALAN on plants by showing that ALAN can significantly affect plant morphology, reproductive timing, and potentially the fitness of plants. We observed both potentially adaptive responses (increased leaf area and altered biomass allocation) and potential negative impacts (reduced reproductive capacity). These findings have important implications for plant populations and ecosystem functioning in the context of global change. As light pollution continues to increase worldwide, the consequences of ALAN on plants may extend beyond individual-level responses and alter population dynamics, community structure, and ecosystem processes. To fully understand the complex direct and indirect consequences of ALAN on plants and ecosystems, future research should include long-term field experiments that track plant demographic responses to ALAN across multiple growing seasons. Such studies will provide valuable insights into the potential cumulative effects of ALAN on plant fitness, population viability, and ecosystem resilience. Additionally, studies examining species-specific ALAN responses could help identify which plant species or functional groups are most vulnerable to light pollution. This information could guide conservation prioritization efforts and inform management strategies to mitigate the adverse effects of ALAN on sensitive plant communities.

## CRediT authorship contribution statement

YL conceived the idea and designed the long-term grassland ALAN manipulation experiment. SW and YL conceived the idea of the present experiment and designed it. SW, ZW, and LX performed the experiment. SW and YL analyzed the data. SW and LX wrote the first draft of the manuscript, with major inputs from YL, and further inputs from AO, ZW, LX, and HZ.

## Acknowledgments

We thank the Erguna Forest-Steppe Ecotone Ecosystem Research Station, Institute of Applied Ecology, Chinese Academy of Sciences for providing the experimental field site and practical assistance with the experiment setup. We thank Xue Zhang, Zhengkuan Lu for their help in the experiment setup.

## Data availability

Data will be made available on request.

## References

Abonyo, C.R.K., Oduor, A.M.O., 2024. Artificial night-time lighting and nutrient enrichment synergistically favor the growth of alien ornamental plant species over co□occurring native plants. J. Ecol. 112(2), 319–337. 10.1111/1365-2745.14235.

Bennie, J., Davies, T.W., Cruse, D., Gaston, K.J., Swenson, N., 2016. Ecological effects of artificial light at night on wild plants. J. Ecol. 104(3), 611–620. 10.1111/1365-2745.12551.

Bennie, J., Davies, T.W., Cruse, D., Inger, R., Gaston, K.J., 2015. Cascading effects of artificial light at night: resource-mediated control of herbivores in a grassland ecosystem. Phil. Trans. R. Soc. B 370(1667), 20140131. 10.1098/rstb.2014.0131.

Bennie, J., Davies, T.W., Cruse, D., Inger, R., Gaston, K.J., Lewis, O., 2018. Artificial light at night causes top□down and bottom□up trophic effects on in vertebrate populations. J. Appl. Ecol. 55(6), 2698–2706. 10.1111/1365-2664.13240.

Bordage, S., Sullivan, S., Laird, J., Millar, A.J., Nimmo, H.G., 2016. Organ specificity in the plant circadian system is explained by different light inputs to the shoot and root clocks. New Phytol. 212(1), 136–149. 10.1111/nph.14024.

Briggs, W.R., 2006. Physiology of plant responses to artificial lighting.

Bürkner, P.-C., 2017. brms: An R Package for Bayesian Multilevel Models Using Stan. J. Stat. Softw. 80(1), 1–28. 10.18637/jss.v080.i01.

Cathey, H.M., Campbell, L.E., 1975. Effectiveness of five vision-lighting sources on photo-regulation of 22 species of ornamental plants. J. Am. Soc. Hort. Sci. 100(1), 65–71.

Cieraad, E., Strange, E., Flink, M., Schrama, M., Spoelstra, K., 2023. Artificial light at night affects plant–herbivore interactions. J. Appl. Ecol. 60(3), 400–410. 10.1111/1365-2664.14336.

Davies, T.W., Smyth, T., 2018. Why artificial light at night should be a focus for global change research in the 21st century. Global Change Biol. 24(3), 87 2-882. 10.1111/gcb.13927.

Diamantopoulou, C., Christoforou, E., Dominoni, D.M., Kaiserli, E., Czyzewski, J. , Mirzai, N., Spatharis, S., 2021. Wavelength-dependent effects of artificial light at night on phytoplankton growth and community structure. Proc. R. Soc. B 288(1953), 20210525. 10.1098/rspb.2021.0525.

Dodd, A.N., Belbin, F.E., Frank, A., Webb, A.A., 2015. Interactions between circadian clocks and photosynthesis for the temporal and spatial coordination of metabolism. Front Plant Sci 6, 245. 10.3389/fpls.2015.00245.

Dorji, T., Totland, O., Moe, S.R., Hopping, K.A., Pan, J., Klein, J.A., 2013. Plant functional traits mediate reproductive phenology and success in response to experimental warming and snow addition in Tibet. Global Change Biol. 19(2), 459–472. 10.1111/gcb.12059.

Falchi, F., Cinzano, P., Duriscoe, D., Kyba, C.C.M., Elvidge, C.D., Baugh, K., Portnov, B.A., Rybnikova, N.A., Furgoni, R., 2016. The new world atlas of artificial night sky brightness. Sci. Adv. 2(6), e1600377. 10.1126/sciadv.1600377.

Famiglietti, C.A., Worden, M., Anderegg, L.D.L., Konings, A.G., 2024. Impacts of climate timescale on the stability of trait-environment relationships. New Phytol. 241(6), 2423–2434. 10.1111/nph.19416.

Fankhauser, C., Staiger, D., 2002. Photoreceptors in Arabidopsis thaliana: light perception, signal transduction and entrainment of the endogenous clock. Planta 216(1), 1–16. 10.1007/s00425-002-0831-4.

Ffrench-Constant, R.H., Somers-Yeates, R., Bennie, J., Economou, T., Hodgson, D., Spalding, A., McGregor, P.K., 2016. Light pollution is associated with earlier tree budburst across the United Kingdom. Proc. R. Soc. B 283(1833), 20160813. 10.1098/rspb.2016.0813.

Gabry, J., Simpson, D., Vehtari, A., Betancourt, M., Gelman, A., 2019. Visualization in Bayesian Workflow. J. R. Stat. Soc. A 182(2), 389–402. 10.1111/rssa.12378.

Gaston, K.J., 2018. Lighting up the nighttime. Science 362(6416), 744–746. 10.1126/science.aau8226.

Gaston, K.J., Bennie, J., Davies, T.W., Hopkins, J., 2013. The ecological impacts of nighttime light pollution: a mechanistic appraisal. Biol. Rev. 88(4), 912–927. 10.1111/brv.12036.

Gaston, K.J., Davies, T.W., Nedelec, S.L., Holt, L.A., 2017. Impacts of Artificial Light at Night on Biological Timings. Annu Rev. Ecol. Evol. Syst. 48(1), 49–68. 10.1146/annurev-ecolsys-110316-022745.

Gaston, K.J., Visser, M.E., Hölker, F., 2015. The biological impacts of artificial light at night: the research challenge. Phil. Trans. R. Soc. B 370(1667), 20 140133. 10.1098/rstb.2014.0133.

Giavi, S., Fontaine, C., Knop, E., 2021. Impact of artificial light at night on diurnal plant-pollinator interactions. Nat. Commun. 12(1), 1690. 10.1038/s41467-021-22011-8.

Gratani, L., 2014. Plant Phenotypic Plasticity in Response to Environmental Factors. Advances in Botany 2014, 1–17. 10.1155/2014/208747.

Grenis, K., Murphy, S.M., 2019. Direct and indirect effects of light pollution on the performance of an herbivorous insect. Insect Sci. 26(4), 770–776. 10.1111/1744-7917.12574.

Gupta, A., Rico-Medina, A., Cano-Delgado, A.I., 2020. The physiology of plant responses to drought. Science 368(6488), 266–269. 10.1126/science.aaz7614.

Haque, M.S., Kjaer, K.H., Rosenqvist, E., Ottosen, C.O., 2015. Continuous light increases growth, daily carbon gain, antioxidants, and alters carbohydrate metabolism in a cultivated and a wild tomato species. Front. Plant Sci. 6, 52 2. 10.3389/fpls.2015.00522.

Heinen, R., 2021. A spotlight on the phytobiome: Plant-mediated interactions in an illuminated world. Basic Appl. Ecol. 57, 146–158. 10.1016/j.baae.2021.10.007.

Hertel, M.F., Araújo, H.H., Stolf-Moreira, R., Pereira, J.D., Pimenta, J.A., Bianchini, E., Oliveira, H.C., 2021. Different leaf traits provide light-acclimation responses in two neotropical woody species. Theoretical and Experimental Plant Physiology 33(4), 313–327. 10.1007/s40626-021-002131.

Hölker, F., Bolliger, J., Davies, T.W., Giavi, S., Jechow, A., Kalinkat, G., Longcore, T., Spoelstra, K., Tidau, S., Visser, M.E., Knop, E., 2021. 11 Pressing Research Questions on How Light Pollution Affects Biodiversity. Front. Ecol. Evol. 9, 767177. 10.3389/fevo.2021.767177.

Hölker, F., Wolter, C., Perkin, E.K., Tockner, K., 2010. Light pollution as a biodiversity threat. Trends Ecol. Evol. 25(12), 681–682. 10.1016/j.tree.2010.09.007.

Irwin, A., 2018. The dark side of light: how artificial lighting is harming the natural world. Nature 553(7688), 268–271. 10.1038/d41586-018-00665-7.

Jiang, H., Shui, Z., Xu, L., Yang, Y., Li, Y., Yuan, X., Shang, J., Asghar, M.A., Wu, X., Yu, L., Liu, C., Yang, W., Sun, X., Du, J., 2020. Gibberellins modulate shade-induced soybean hypocotyl elongation downstream of the mutual promotion of auxin and brassinosteroids. Plant Physiol. Biochem. 15 0, 209–221. 10.1016/j.plaphy.2020.02.042.

Jiang, Y., Deng, Y., Yang, W., Dang, R., 1996. Anatomic studies of the floral nectaries in Elsholtzia densa Benth. Acta Bot. Boreali-Occidential Sinica 16(3), 239–244.

Kim, Y.J., Yu, D.J., Rho, H., Runkle, E.S., Lee, H.J., Kim, K.S., 2015. Photosynthetic changes in Cymbidium orchids grown under different intensities of night interruption lighting. Sci. Hortic. 186, 124–128. 10.1016/j.scienta.2015.01.036.

Knop, E., Zoller, L., Ryser, R., gerpe, C., Hörler, M., Fontaine, C., 2017. Artificial light at night as a new threat to pollination. Nature 548(7666), 206–209. 10.1038/nature23288.

Krittika, S., Yadav, P., 2019. Circadian clocks: an overview on its adaptive significance. Biol. Rhythm Res. 51(7), 1109–1132. 10.1080/09291016.2019.1581480.

Kudo, G., Harder, L.D., 2005. Floral and inflorescence effects on variation in pollen removal and seed production among six legume species. Funct. Ecol. 19(2), 245–254. 10.1111/j.1365-2435.2005.00961.x.

Kwak, M., Je, S., Cheng, H., Seo, S., Park, J., Baek, S., Khaine, I., Lee, T., Jang, J., Li, Y., Kim, H., Lee, J., Kim, J., Woo, S., 2018. Night Light-Adaptation Strategies for Photosynthetic Apparatus in Yellow-Poplar (Liriodendron tulipifera L.) Exposed to Artificial Night Lighting. Forests 9(2), 74. 10.3390/f9020074.

Kwak, M.J., Lee, S.H., Khaine, I., Je, S.M., Lee, T.Y., You, H.N., Lee, H.K., Jan g, J.H., Kim, I., Woo, S.Y., 2017. Stomatal movements depend on interactions between external night light cue and internal signals activated by rhythmic starch turnover and abscisic acid (ABA) levels at dawn and dusk. Ac ta Physiol. Plant. 39(8), 162. 10.1007/s11738-017-2465-y.

Kyba, C.C.M., Kuester, T., Miguel, A.S.d., Baugh, K., Jechow, A., Hölker, F., Bennie, J., Elvidge, C.D., Gaston, K.J., Guanter, L., 2017. Artificially lit surface of Earth at night increasing in radiance and extent. Sci. Adv. 3(11), e1701528. 10.1126/sciadv.1701528.

Lazzarin, M., Meisenburg, M., Meijer, D., van Ieperen, W., Marcelis, L.F.M., Kappers, I.F., van der Krol, A.R., van Loon, J.J.A., Dicke, M., 2021. LEDs Make It Resilient: Effects on Plant Growth and Defense. Trends Plant Sci. 26(5), 496–508. 10.1016/j.tplants.2020.11.013.

Li, F., Chen, B., Xu, K., Gao, G., Yan, G., Qiao, J., Li, J., Li, H., Li, L., Xiao, X., Zhang, T., Nishio, T., Wu, X., 2016. A genome-wide association study of plant height and primary branch number in rapeseed (Brassica napus). Plant Sci. 242, 169–177. 10.1016/j.plantsci.2015.05.012.

Liao, Y., Suzuki, K., Yu, W., Zhuang, D., Takai, Y., Ogasawara, R., Shimazu, T., Fukui, H., 2014. Night break effect of LED light with different wavelengths on floral bud differentiation of Chrysanthemum morifolium Ramat ’Jim ba’and ’Iwa’ no hakusen. Environ. Control Biol. 52(1), 45–50. 10.2525/ecb.52.45.

Liu, H., Wang, R., Lu, X.T., Cai, J., Feng, X., Yang, G., Li, H., Zhang, Y., Han, X., Jiang, Y., 2021. Effects of nitrogen addition on plant-soil micronutrients vary with nitrogen form and mowing management in a meadow steppe. Environ. Pollut. 289, 117969. 10.1016/j.envpol.2021.117969.

Lockett, M.T., Rasmussen, R., Arndt, S.K., Hopkins, G.R., Jones, T.M., 2022. Artificial light at night promotes bottom-up changes in a woodland food chain. Environ. Pollut. 310, 119803. 10.1016/j.envpol.2022.119803.

Macgregor, C.J., Evans, D.M., Fox, R., Pocock, M.J., 2017. The dark side of street lighting: impacts on moths and evidence for the disruption of nocturnal pollen transport. Global Change Biol. 23(2), 697–707. 10.1111/gcb.13371.

MacGregor, C.J., Pocock, M.J., Fox, R., Evans, D.M., 2015. Pollination by nocturnal Lepidoptera, and the effects of light pollution: a review. Ecol. Entomol. 40(3), 187–198. 10.1111/een.12174.

Meng, L., Zhou, Y., Román, M.O., Stokes, E.C., Wang, Z., Asrar, G.R., Mao, J., Richardson, A.D., Gu, L., Wang, Y., 2022. Artificial light at night: an under-appreciated effect on phenology of deciduous woody plants. PNAS nexus 1(2), pgac046. 10.1093/pnasnexus/pgac046/6569705.

Meravi, N., Kumar Prajapati, S., 2020. Effect street light pollution on the photosynthetic efficiency of different plants. Biol. Rhythm Res. 51(1), 67–75. 10.1080/09291016.2018.1518206.

Patke, A., Young, M.W., Axelrod, S., 2020. Molecular mechanisms and physiological importance of circadian rhythms. Nat. Rev. Mol. Cell Biol. 21(2), 67–84. 10.1038/s41580-019-0179-2.

Pierik, R., de Wit, M., 2014. Shade avoidance: phytochrome signalling and other aboveground neighbour detection cues. J. Exp. Bot. 65(11), 2815–2824. 10.1093/jxb/ert389.

Poorter, H., Niinemets, U., Poorter, L., Wright, I.J., Villar, R., 2009. Causes and consequences of variation in leaf mass per area (LMA): a meta-analysis. New Phytol. 182(3), 565–588. 10.1111/j.1469-8137.2009.02830.x.

Quail, P.H., 2002. Photosensory perception and signalling in plant cells: new paradigms? Curr. Opin. Cell Biol. 14(2), 180–188. 10.1016/s0955-0674(02)00309-5.

R Core Team, 2020. R: A language and environment for statistical computing. R Foundation for Statistical Computing, Vienna, Austria.

Rodger, J.G., Bennett, J.M., Razanajatovo, M., Knight, T.M., van Kleunen, M., Ashman, T.-L., Steets, J.A., Hui, C., Arceo-Gómez, G., Burd, M., 2021. Widespread vulnerability of flowering plant seed production to pollinator declines. Sci. Adv. 7(42), eabd3524. 10.1126/sciadv.abd3524.

Sanders, D., Frago, E., Kehoe, R., Patterson, C., Gaston, K.J., 2021. A meta-analysis of biological impacts of artificial light at night. Nat. Ecol. Evol. 5(1), 74–81. 10.1038/s41559-020-01322-x.

Schupp, E.W., Jordano, P., Gomez, J.M., 2010. Seed dispersal effectiveness revisited: a conceptual review. New Phytol. 188(2), 333–353. 10.1111/j.1469-8137.2010.03402.x.

Secondi, J., Davranche, A., Théry, M., Mondy, N., Lengagne, T., Isaac, N., 2019. Assessing the effects of artificial light at night on biodiversity across latitude – Current knowledge gaps. Global Ecol. Biogeogr. 29(3), 404–419. 10.1111/geb.13037.

Segrestin, J., Mondy, N., Boisselet, C., Guigard, L., Lengagne, T., Poussineau, S., Secondi, J., Puijalon, S., 2021. Effects of artificial light at night on the leaf functional traits of freshwater plants. Freshwat. Biol. 66(12), 2264–2271. 10.1111/fwb.13830.

Sena, S., Kumari, S., Kumar, V., Husen, A., 2024. Light emitting diode (LED) lights for the improvement of plant performance and production: A comprehensive review. Current Res. Biotechnol. 7, 100184. 10.1016/j.crbiot.2024.100184.

Singhal, R.K., Kumar, M., Bose, B., 2019. Eco-physiological Responses of Artificial Night Light Pollution in Plants. Russ. J. Plant Physiol. 66(2), 190–202. 10.1134/s1021443719020134.

Speißer, B., Liu, Y., van Kleunen, M., Oduor, A., 2021. Biomass responses of widely and less□widely naturalized alien plants to artificial light at night. J. Ecol. 109(4), 1819–1827. 10.1111/1365-2745.13607.

Vehtari, A., Gelman, A., Gabry, J., 2017. Practical Bayesian model evaluation using leave-one-out cross-validation and WAIC. Statistics and Computing 27, 1413–1432. 10.1007/s11222-016-9696-4.

Violle, C., Navas, M.-L., Vile, D., Kazakou, E., Fortunel, C., Hummel, I., Garnier, E., 2007. Let the concept of trait be functional! Oikos 116(5), 882–892. 10.1111/j.0030-1299.2007.15559.x.

Waser, N.M., Ollerton, J., 2006. Plant-pollinator interactions: from specialization to generalization. University of Chicago Press.

Webb, A.A., 2003. The physiology of circadian rhythms in plants. New Phytol. 1 60(2), 281-303. 10.1046/j.1469-8137.2003.00895.x.

Yerushalmi, S., Green, R.M., 2009. Evidence for the adaptive significance of circ adian rhythms. Ecol. Lett. 12(9), 970–981. 10.1111/j.1461-0248.2009.01343.x.

Zhao, K., Ma, B., Xu, Y., Stirling, E., Xu, J., 2021. Light exposure mediates circ adian rhythms of rhizosphere microbial communities. ISME J. 15(9), 2655–2664. 10.1038/s41396-021-00957-3.

Zheng, Q., Teo, H.C., Koh, L.P., 2021. Artificial Light at Night Advances Spring Phenology in the United States. Remote Sens. 13(3), 399. 10.3390/rs13030399.

